# The Importance of Synchrony in the Neural Control of Movement

**DOI:** 10.64898/2026.06.26.734805

**Authors:** Masashi Hasegawa, Barbara Gruszka, Michael S. Finch, Vikshar J. Athreya, Aaron D. Milstein, Ian Antón Oldenburg

## Abstract

Throughout the mammalian cortex, populations of neurons must work together to enact behaviors. While population recordings in motor cortex have revealed many aspects of when neurons fire during behaviors, limitations in causal experiments have made it difficult to identify which features of neural activity directly drive movements and which do not. Here, we explicitly test the principles of neural coding using high temporal precision multiphoton holographic optogenetics in the motor cortex. We show that activation of a small number (50-75) of Layer 2/3 excitatory neurons in the motor cortex is sufficient to drive movements. The efficacy of stimulation-driven movement depends on the state of the local circuit, and, to a lesser extent, the identity of which neurons are stimulated. We test whether evoked activity is acting through a ‘rate code’ or a ‘timing code’ by holding the firing rate and cell identities constant while varying the millisecond precise timing of activation. We find an unexpected and strong dependence on inter-cell synchrony when evoking movements. This neural synchrony recruits distinct patterns of recurrent excitation and inhibition. These findings provide evidence that the timing code, more so than the rate code, drives motor output.

## Main

Even simple actions involve the coordinated activity of many neurons across motor-related brain regions. Over the last few decades, technological advances have enabled observing the activity of large populations of neurons^1–4^, leading to sophisticated ‘population code’ models of how these neurons might work together to drive behaviors^5–7^. While these approaches allow us to sample from hundreds or thousands of neurons simultaneously, our ability to causally manipulate population activity is far more limited, typically restricted to driving functionally or genetically identified cell types uniformly^8–11^. This leads to a fundamental mismatch in our ability to test which features of neural codes are involved in evoking behaviors, and which features are not.

We focus on a long-standing debate on the nature of neural activity in cortex^12–14^: whether relevant activity is encoded in the number of action potentials a neuron fires, i.e. a ‘rate code’, or in the millisecond precise timing of these action potentials, i.e. a ‘timing code’. The rate-based models of neural population activity are able to describe sensory stimuli^15–18^, motor behaviors^19,20^, and predict future actions^21–25^, demonstrating that rate-based analyses are able to reveal important features of neural activity. At the same time, individual cortical neurons are observed to fire action potentials with millisecond precision, and can exhibit different responses based on small changes in the timing of excitatory and inhibitory inputs^26–32^. While precise temporal codes are well described in songbirds^33^, and rodent subcortical structures^34^; their role in mammalian cortex remains elusive with evidence both for^35,36^ and against^37,38^. In the motor cortex, time varying neural dynamics with timescales on the order of tens to hundreds of milliseconds have been correlated to behaviors^38,39^, but whether this effect extends to smaller timescales, and whether these features causally drive the behaviors remains unknown.

To directly test the role of millisecond timing in population codes, we use high temporal precision multiphoton holographic optogenetics^40,41^ in the mouse caudal forelimb area (CFA)^42^, precisely activating small numbers of cells. While many forms of multiphoton optogenetics allow selective activation of cells with cellular resolution^43–69^, temporally focused, expanded spot holographic activation, such as 3D-SHOT in combination with a fast opsin, also allows millisecond precise control of the timing of evoked action potentials, with a low rate of failures or doublets^62–69^. When paired with fast spatial light modulators (SLMs), this combination of approaches allows for exceptional control over the timing of activity across multiple neurons. This allows us to experimentally control which cells we are activating, the degree to which we drive each cell, and their precise inter-cell synchrony.

We find that relatively few neurons are needed to evoke a movement in a naïve animal. Among sets of 75 co-stimulated Layer 2/3 (L2/3) motor cortex neurons, more than half evoked movements, with ongoing brain state playing a key role in gating the initiation of movements. Notably, a neuron’s activity in response to a spontaneous movement was a poor predictor of whether it would evoke a movement, highlighting motor cortex’s multifunctionality in both representing and enacting movements. To test the relative importance of rate and timing codes, we designed multiphoton optogenetic stimulation patterns where the same neurons were each driven at the same rate but differed in the inter-cell synchrony of activity. We find that the main determinant of the effectiveness of stimulation is the inter-cell timing, with asynchronous stimulation failing to drive movements. While all stimuli drive recurrent inhibition, recurrent excitation is more sensitive to synchrony. Our data shows that both rate and timing play a role in evoking movements, but the timing features are more pronounced and follow a shorter timescale than is commonly analyzed.

## Holographic stimulation evokes body movements

While electrical^42,70–73^ or optogenetic stimuli^74,75^ in motor cortex can evoke across species, these stimuli typically activate hundreds of neurons non-specifically. We first sought to determine the number of Layer 2/3 (L2/3) excitatory neurons sufficient to evoke a movement in an awake naïve mouse. We performed scanless multiphoton holographic optogenetic stimulation while simultaneously imaging *in vivo* calcium activity in motor cortex (Fig. 1a). We optimized our holographic system to ensure our stimulation resolution is approximately the size of a cortical pyramidal cell (17.1 ± 0. 9 µm radial, 29.6 ± 2.3 µm axial (n = 40 cells), Fig.1b). As done previously^63,67,68^, each targeted cell was calibrated to receive a bespoke power, and stimulated cells responded reliably, even in large groups (Fig. 1c). We then stimulated ensembles of randomly selected photoactivatable cells in naïve animals and monitored forelimb movements with a high-speed camera (Fig. 1c, d). Cortical activity in the motor cortex is heterogeneous, but can change and become more stereotyped over learning^76^. To ensure we studied endogenous network states and avoid triggering over trained behaviors, we stimulated neuronal ensembles in naïve mice in the absence of a behavioral task.

**Fig. 1.**
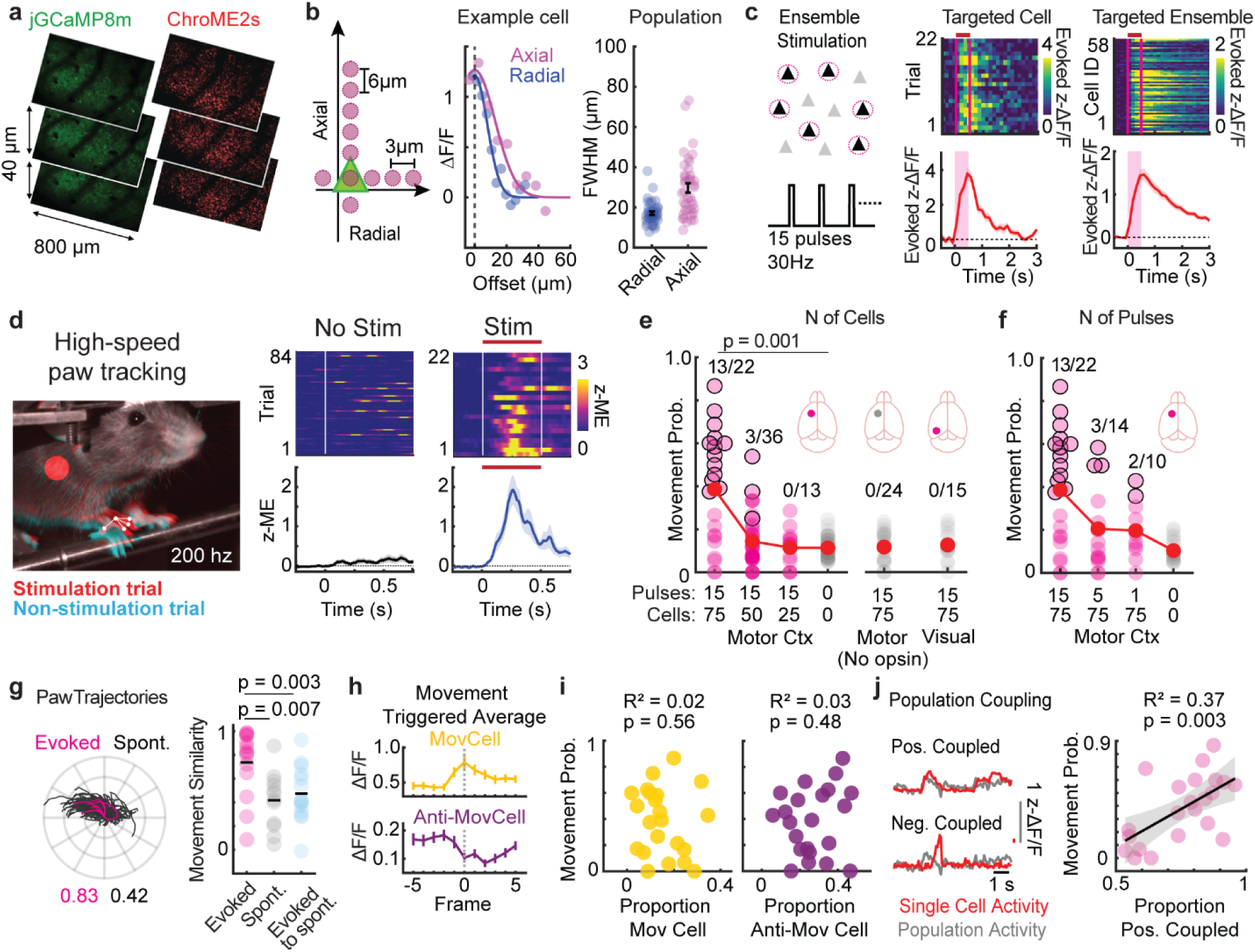
Two-photon holographic stimulation of the motor cortex evokes body movements. **a**. Volumetric 2p-images of Layer 2/3 motor cortex with jGCaMP8m (left) and ChroME2s-p2a-mRuby3 (right) in excitatory neurons. **b**. Schematic of in vivo physiological point spread function (PPSF) experiment (left). Example PPSF (middle), measured ΔF/F with radial (blue) and axial (maroon) offsets. Solid line gaussian fit. Right: measured PPSF widths (n = 40 cells, 4 FOVs, 2 mice). **c**. Schematic of *in vivo* two-photon holographic stimulation, 15 pulses at 30 Hz. Right: Neural response of an example neuron over trials and each member of a targeted ensemble. **d**. Left: two frames superimposed from high-speed (200 Hz) paw tracking. Red: stim trial, Cyan: no stim trial. Right: z-scored motion energy (z-ME) from a ROI including the right paw with and without activation of 75 neurons. **e-f**. Evoked movement probability across ensembles with different conditions. Red circles are group mean. Inset number and black outline represent ensembles that were statistically different from their paired no stim control (Fisher Exact Test). 75 cells with 15 pulses was significantly higher than the no stim condition (p=0.001, signed rank test). Number of cells experiment: n = 71 ensembles, 29 FOVs, 10 mice; No opsin control in motor cortex: n = 24 ensembles, 8 FOVs, 3 mice; Opsin in visual cortex control: n = 15 ensembles, FOV = 6, 3 mice; Number of pulses experiment: n = 46 ensembles, 19 FOVs, 9 mice. **g**. Example paw trajectories evoked (magenta) and spontaneous (black), cosine similarity of vector below. Cosine similarity of all movements (right) (n = 14 ensembles, 12 FOVs, 8 mice from 75 cell ensembles stimulation). **h**. Motion triggered average activity of example movement correlated and anticorrelated cells. **i**. Movement probability as a function of the proportions of movement-correlated/anti-correlated cells in each ensemble. **j**. Left: Example positively and negatively coupled cells. Right: Movement probability as a function of the proportion of positive population-coupled cells in each ensemble.

We found that simultaneous activation of 50 or 75 neurons, driven with 15 pulses at 30 Hz, evoked paw movements (13/22 and 3/36 ensembles evoked movements for 75 and 50 neurons, Fisher exact test, Fig.1d-f). The probability of evoking a movement increased with both the size of the ensemble and the number of pulses (Fig.1e, f, Extended Data Fig. 1a, b). Neither stimulation in the visual cortex nor in opsin-negative animals elicited body movements in any tested ensemble (0/15, and 0/24 ensembles in visual cortex and opsin-negative motor cortex, respectively, evoked movement Fig.1e, Extended Data Fig. 1c, d). Furthermore, stimulation that occurred during or immediately after an ongoing movement rarely evoked an additional movement (Extended Data Fig. 1e-f).

High speed paw tracking revealed that evoked movements were typically stereotyped (Fig. 1g, Extended Data Fig. 1g, cosine similarity for ensemble stimulation 0.74 ± 0.07, spontaneous 0.42 ± 0.07, *p* = 0.007, ensembles vs spontaneous 0.47 ± 0.06, *p* = 0.003), although a wide variety of movements were observed across ensembles and mice.

Across all tested ensembles, the probability of evoking a movement was heterogenous, ranging from 0 to 86.6% of trials (Fig. 1d-f). Movement probability had a weak negative correlation with the amplitude of stimulation-evoked neural response in targeted ensembles, indicating stimulation failures were not responsible for heterogeneity (R^2^ = 0.19, *p* = 0.04, Extended Data Fig. 1h). Similarly, neither the spatial placement of targeted cells within ensembles (Target cell spread vs Evoked responses, R^2^ = 0.01, *p* = 0.70, or vs Movement probability, R^2^ = 0.03, *p* = 0.44, Extended Data Fig. 1i), nor their observed activity level during non-stimulation trials predicted their efficacy (Movement probability vs proportion of high activity cells in a ensemble, R^2^ = 0.04, *p* = 0.37, Movement probability vs proportion of low activity cells, R^2^ = 0.00, *p* = 0.96, Extended Data Fig. 1j).

Next, we hypothesized that motor cortex ‘movement neurons’ (i.e., neurons that fire during spontaneous movements, Fig. 1h) would be involved in driving movements; and thus, targeting them would be more effective at driving evoked movements. By calculating movement-triggered averages during non-stimulation trials, we categorized cells as either movement-active, movement-suppressed, or neither (see Methods). However, the proportion of cells within an ensemble belonging to either category was not predictive of evoked movement probability (Movement probability vs Proportion of movement-cells in a ensemble, R^2^ = 0.02, *p* = 0.56, Movement probability vs anti-movement cells, R^2^ = 0.03, *p* = 0.48, Fig. 1i).

Alternatively, we reasoned that the relationship between a cell and its local network may gate its efficacy in evoking movements. To quantify this relationship, we calculated population coupling^77^ (see Methods). Cells with high coupling track the population activity closely, whereas minimally coupled cells fire independently of the local environment. We found that ensembles with high proportion of positively coupled cells were associated with higher movement probability (Fig. 1j, R^2^ = 0.37, *p* = 0.003), indicating that cells embedded in local population dynamics are preferentially involved in initiating movements in the motor cortex.

Overall, our data show that holographically evoked movements depend on the number of targeted cells and the number of evoked action potentials, and only weakly on the identity of the targeted neurons.

## Ongoing neural activity predicts the behavioral effect of holographic stimulation

Given the relatively weak impact of cell identity on the generation of movements, we sought to explore how ongoing neural activity in motor cortex may permit or block holographic stimulation from eliciting a movement, i.e. is there a ‘permissive brain state’ observable immediately preceding stimulation that predicts whether stimulation will result in a movement. We focused on activity before stimulation to avoid the confound of stimulation-evoked activity influencing the observed activity.

We designed several ensemble-based linear models that related the activity of neurons in the 5 frames (0.67s) before stimulation, to the movement energy evoked by that ensemble (see Methods). As our experiments have relatively few stimulation trials for the number of recorded neurons, models that assigned weights to each neuron easily overfit, whereas those that assigned weights to activity in a small number of principal components extracted from non-stimulation trials (see Methods) failed to fit (All neuron model: ΔR^2^ _model – shuffle_ = 1.7e-6 ± 7.1e-7; Top four principle components model: ΔR^2^ _model – shuffle_ = -0.07 ±0.04. Extended Data Fig. 2).

We therefore devised a modeling approach that allowed us to combine neural activity across datasets using interpretable features. We categorized cells based on two criteria: (1) whether they were targeted for photostimulation, and (2) how correlated they were to evoked movements (i.e. the top 5% most correlated, top 5% most anti correlated, and all other cells) (Fig. 2a, Extended Data Fig. 3a-c). The mean activity of each of these six groups of neurons in the 5 frames prior to stimulation was used to predict stimulation-evoked motion using linear regression model (see Methods). The model predicted the true evoked motion energy much better than a shuffle control (Fig. 2b-c, ΔR^2^_model – shuffle_ = 0.41±0.12), and was cross validated using either held out trials, held out ensembles, or by shuffling the correlation groups independently or yoked to the shuffled data (Extended Data Fig. 3d-h).

**Fig. 2.**
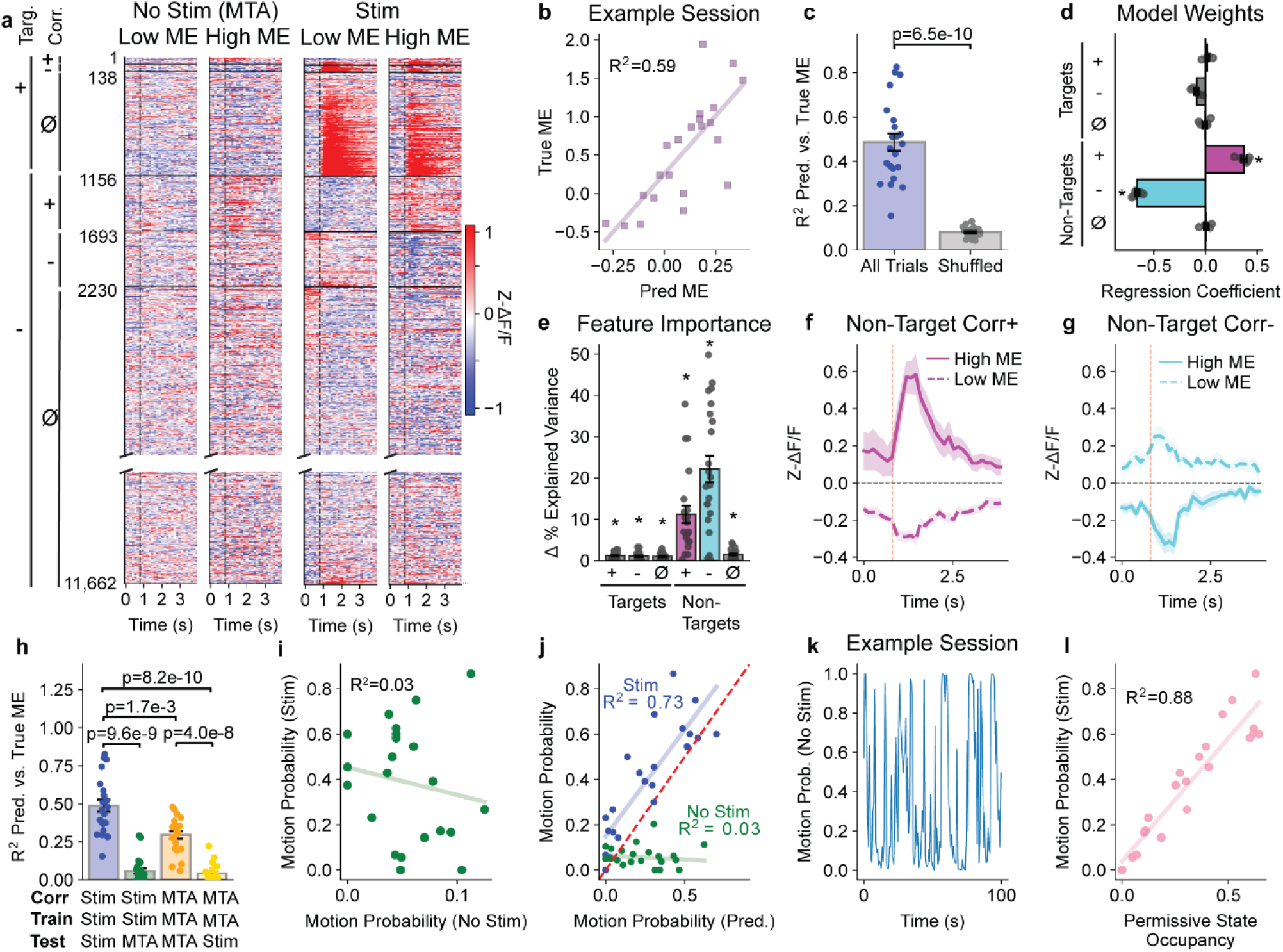
Neural activity before stimulation predicts evoked movement rate. **a**. Mean response (z-scored dF/F) of recorded neurons aligned to spontaneous movements (movement triggered average, MTA) in no stim trials (left) or stimulation onset in stim trials (right). Trials are divided into high or low motion energy (ME). Cells are divided into six categories based on whether they are in a stimulation targeted ensemble (Targ.), and whether their activity was correlated positively (+), negatively (-), or uncorrelated (*ϕ*) with evoked ME (Corr.). **b**. Pre-stimulation activity predicts evoked movement. Linear regression was used to predict each stim trial’s evoked ME from the mean activity of each group in a during the first 5 frames before stimulation. For each trial from an example ensemble, evoked ME is compared to predicted ME (R^2^=0.59, p=3.3e-5, Pearson). **c**. Model explained variance (Pearson R^2^) for each ensemble vs the average of 200 models with ME values shuffled for all stim trials with the same ensemble. **d**. Model coefficients across five cross-validations with held-out-trials. **e**. Parameter importance: the change in explained variance when single neural features are removed during model training. **f-g**. Trial- and cell-averaged activity of the most predictive groups: non-targeted positive or negatively correlated cells, separated by trials with high and low evoked ME. **h**. When models are trained on data from stim trials, using neural features based on movement correlations from stim trials (corr: stim), they fail to predict spontaneous movements (test: stim vs MTA, p=9.6e-9, blue vs green). When models are trained on data from no stim trials with features based on movement correlations from no stim trials (corr. and train: MTA), they fail to predict stim-evoked movements (test: MTA vs stim, p =4.0e-8, orange vs yellow). **i**. The fraction of high ME no stim trials does not predict the fraction of high ME stim trials (R^2^=0.03, p=0.47). **j**. Binary classifiers were trained with logistic regression on data from stim trials to predict either the fraction of high ME stim trials (blue, R^2^=0.73, p=4.3e-7), or the fraction of high ME no stim trials (green, R^2^=0.03, p=0.43). **k**. For an example no stim trial, the model trained on data from stim trials was used to predict the likelihood that stimulation would have produced high ME (permissive state) in successive 5 frame windows. **l**. The fraction of time spent in the permissive state during no stim trials predicts the observed fraction of high ME stim trials for each ensemble (R^2^=0.88, p=1.2e-10, Pearson).

We further validated our model by using neural activity within the same post-stimulation window as the motion energy. This ‘post’ model conflates movement activity with optogenetically evoked activity but provides an upper bound for how accurate this model could be (Extended Data Fig. 3i-l, R^2^=0.89±0.01, Shuffled R^2^=0.06±0.002). Taken together, we conclude that the ongoing neural activity preceding photostimulation can predict the probability and magnitude of evoked movement.

We then sought to determine which cell classes contribute to the accurate prediction of stimulation-evoked movement. Surprisingly, we find that pre-stimulus activity of directly targeted cells has little effect on the model, both being assigned low weights and their removal from the model resulting in little change in accuracy (Fig. 2d-e). In contrast, the model consistently weights and relies upon untargeted cells that are most correlated, either positively or negatively, with movements (Fig. 2d-g). These cell groups show clear divergence in their pre-stimulus activity between trials with high vs. low evoked movement energy (Fig. 2 f-g). We thus conclude that ongoing network activity, and not merely the activity of directly targeted cells, determines whether the brain is in a state that is permissive for stimulus-evoked movements.

To distinguish if this pre-stimulus state was a ‘permissive state’, i.e. requires stimulation to evoke a movement, or merely a ‘predictive state’, i.e. indicates a movement is about to happen, we examined how models trained to predict stimulation-evoked motion performed in predicting motion energy on non-stimulated trials. First, we aligned non-stimulated trials to the largest movement in that trial. Since not all trials contained large movements, the resulting spontaneous motion energy values spanned a large range across trials (Extended Data Fig. 3a-c). The model trained on stimulation-evoked movements failed to accurately predict spontaneous movements (R^2^ _stim pred no stim_=0.06±0.02, Fig. 2h, green). Next, we performed the inverse analysis by training a model to predict spontaneous movement energy on non-stimulated trials from the neural activity just prior to the spontaneous movement (R^2^ _no stim trials_=0.30±0.02, Fig. 2h, orange). This model was unable to predict evoked movement energy on stimulated trials (R^2^ _no stim pred stim_=0.04±0.01, Fig. 2h, yellow). Across ensembles, the fraction of non-stimulated trials with high motion energy was not correlated with the fraction of stimulated trials with high motion energy (Fig. 2i).

We next used the predicted motion energies from the linear regression model to train binary logistic classifiers for each targeted ensemble to discriminate trials that were above or below a threshold for evoked movements (Methods). Accordingly, across all ensembles, the classifier trained to predict evoked movements on stim trials accurately predicted the fraction of trials with high motion energy (R^2^_stim_=0.73) but poorly predicted the fraction of non-stimulated trials with high motion energy (R^2^_no stim_=0.03, Fig. 2j). These results suggest that there is a ‘permissive state’ of cortex that influences whether an external stimulus will evoke a movement, and it is distinct from a ‘predictive’ or ‘preparatory state’ that normally precedes spontaneous movements.

Finally, to determine if this permissive state can explain the heterogeneity in evoked movement probability by different ensembles, we calculated the fraction of time during non-stimulated trials time that the classifier predicts would have resulted in a stimulus-evoked movement had a stimulation occurred (see Methods). Non-stimulated trials naturally oscillated between high and low predicted motion probabilities (Fig. 2k). This approach revealed that the fraction of time the cortex spent in the permissive state within non-stimulated trials correlated well with the fraction of stimulated trials that resulted in evoked movement (R^2^ =0.88, Fig. 2l).

Taken together we conclude that ongoing neural activity before stimulation defines a permissive state that permits evoked movements. Much of the heterogeneity between trials and ensembles is due to the variability in an animal’s neural state, rather than features of the stimulus itself.

## Synchrony of neural activity is critical to drive movements

Having established an effective framework to study cortically evoked movement, we designed an experiment to directly test the relative importance of both rate and timing codes. We compared two distinct evoked activity patterns where in both cases the same cells were driven in the same way, but the two stimuli differed in their intercell timing. In ‘sync’ trials, we activated ensembles of 50-75 cells with 15 pulses at 30Hz as above (Fig. 1c), but in ‘async’ trials we split the ensemble into three interleaved groups, each offset by 5ms (Fig. 3a). In both cases, the same cells were activated with the same 15 pulses at 30Hz, but the population synchrony was altered. Using stroboscopic imaging (Extended Data Fig. 4a, see Methods) we confirm that the system can switch holograms in less than 3 milliseconds (2.13 ms ± 0.09, Fig. 3b, Extended Data Fig. 4a-b). By waiting 5 ms before stimulation, we ensure that each target receives comparable power regardless of its order of stimulation sequence in the async condition (Extended Data Fig. 4c, d). Furthermore, we confirmed that the relative stimulation power delivered to each individual target was not affected by the number of targets included in the hologram (Extended Data Fig. 4e, f). Thus targeted cells receive the same amount of stimulation, regardless of whether they are driven in the ‘sync’ or ‘async’ trial type.

**Fig. 3.**
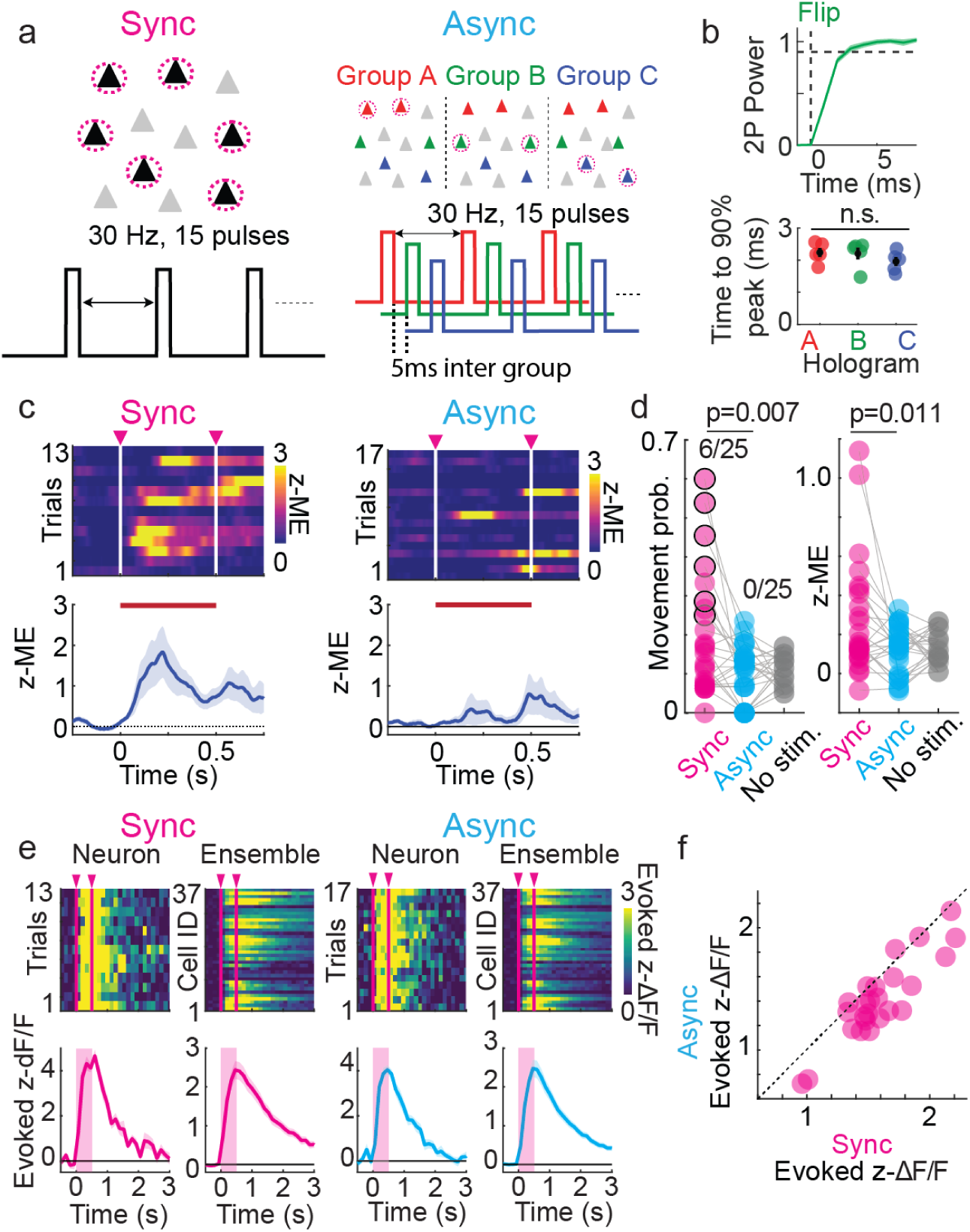
Inter-cell synchrony of neural activation is critical for evoking movements. **a**. Left: Schematic of synchronous and asynchronous activation. In async activation, targeted neurons were randomly separated into three groups and stimulated in an interleaved manner (with 5ms between pulses). The same neurons were stimulated with the same number of stimulation pulses (15 pulses) at 30 Hz. **b**. Two-photon power efficiency (2P-power) during the hologram flipping. Top: Example 2P power transition during the flipping of one hologram. Bottom: Summary of hologram flipping speed. 2P power efficiencies of flipped holograms reached 90% of peak power within 3 ms (p = 0.23, Kruskal-Wallis test, non-significant). **c**. Example movement data (z-ME) in sync and async condition. **d**. Left: Movement probability in sync and async conditions (N = 25 paired holograms, 11 FOVs, 5 mice). 6/25 sync vs 0/25 async holograms significantly evoked paw movements (Fisher Exact Test, inset and black circles), The sync evoked movement probability was higher than async condition (*p* = 0.007, signed-rank test). Right: Mean of the amplitude of movements (Sync vs Async p = 0.011, signed-rank test). **e**. Example neural responses of a single targeted neuron and ensemble in sync (Left) and async condition (Right). **f**. Evoked neural responses of targeted ensembles in sync and async condition. In figure panels, red and white lines on heatmaps represent stimulation onset and offset. Traces shown in b, c, e are mean ± s.e.m.

Remarkably, despite each cell individually receiving the same illumination pattern in both cases, ‘sync’ activation of either 50 or 75 cell ensembles evoked paw movements while ‘async’ activation never evoked movements (6/25 sync- and 0/25 async-evoked movements, Fisher exact test. Movement probability sync vs async, p = 0.007, signed-rank test, Fig. 3c-d, Extended Data Fig. 4g). This result did not arise from a failure of holographic stimulation, because sync and async stimulations activated targeted neurons with a similar cell to cell and trial to trial reliability (Fig. 3e). However, we observed that the amplitude of GCaMP response in targeted cells was slightly but significantly larger (Δz-ΔF/F, 0.178 ± 0.03, signed-rank test, *p* = 9.0e-5) in the sync condition than the async condition. The difference in evoked responses cannot explain the difference in behavior for two reasons: first, the difference in evoked responses between sync to async difference of conditions is smaller than the ensemble-to-ensemble variance (Fig 3f, Δz-ΔF/F = 0.178 between sync and async, SD = 0.31 z-ΔF/F sync and 0.33 z-ΔF/F async) and second, targeted cell activation level is anti-predictive of movement probability (Extended Data Fig. 1h). Instead, these data raise the possibility that synchronous and asynchronous activation engages the local circuit in different ways.

## Synchronous and Asynchronous Stimulation Differentially Recruit Recurrent Excitation and Inhibition

Between sync and async stimulation, each cell individually experienced the same pattern of activation, and each pulse of light was the same duration and intensity (Extended Data Fig. 4a-f). Therefore, we reasoned that the subtle differences in targeted cells’ responses could be the result of the timing dependence of local recurrent circuits. To explore the mechanism of these recurrent circuit effects, we designed an experiment to compare the effects of changing the number of stimulated cells – adding cells synchronously or asynchronously. In addition to the sync and async condition described above we created a ‘subset’ condition, where we illuminate targets in the same way as in the async condition, but only one of the three groups were stimulated (Fig. 4a-b). Thus, we can compare the response in cells, when we activate that subset alone (‘subset’), as part of a larger synchronous group (‘sync’), or as part of a larger asynchronous series (‘async’) (Fig. 4, Extended Data Fig. 5). We restrict our analysis to low movement trials to avoid confounds.

**Fig. 4.**
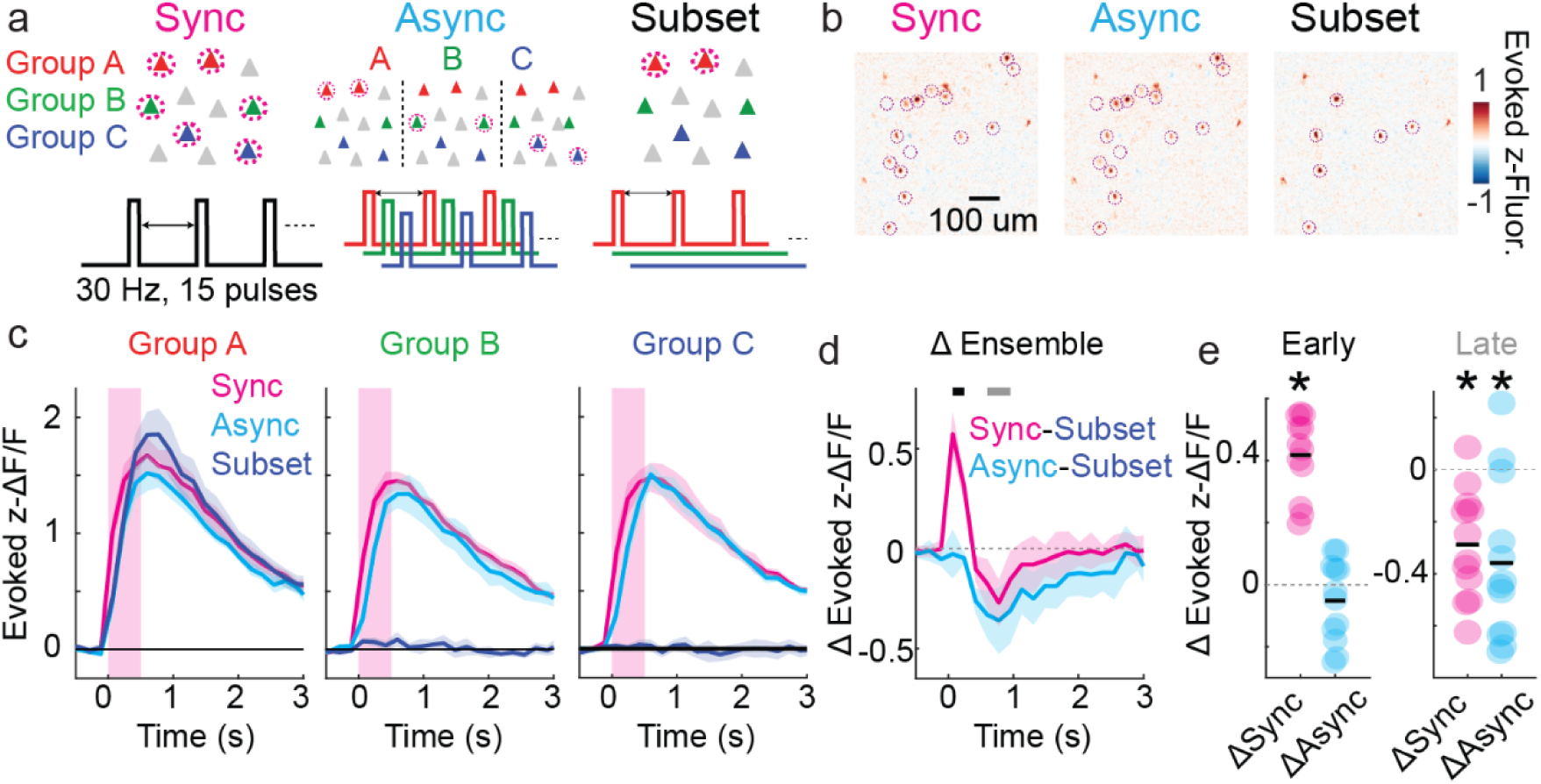
Synchronous activation produced fast rise of neural activity in the targeted ensembles. **a.** Schematics of sync, async and single-subset activations. As above, async condition splits full ensemble into three equal groups, in a single-subset activation, only one of the three groups was stimulated. **b**. Example activation maps of sync, async and single-subset activations in a single session. One plane of four shown. **c**. Mean of targeted cells in each subgroup in response to sync, async and single-subset activation (N = 12 ensembles, 4 FOVs, 2 mice). **d.** Difference in the evoked neural responses between three activations. Black and grey lines indicate the time points analyzed in e. **e**. Difference in the evoked neural responses in early and late time points (Early time point, *p* = 0.0005 for synchronous vs single-subset, *p* = 0.30 for asynchronous vs Single-subset response; Late time point, *p* = 0.0015 for synchronous vs single-subset, *p* = 0.005 for asynchronous vs Single-subset response). Asterisk indicates the statistical differences.

We found that the subset stimulation evoked larger responses in stimulated cells than either the sync or async condition (Fig. 4c), consistent with previous observations that cortical activation drives recurrent inhibition proportional to the number of cells driven^45,68,78^ (Fig. 4d-e ‘late’ activation -0.29 ± 0.06 z-ΔF/F, *p* = 0.0015 for sync-subset, -0.36 ± 0.09 z-ΔF/F, *p* = 0.005 for async-subset, signed-rank test). However, we found that sync activation was able to evoke neural response faster than the async or subset condition (Fig. 4d-e early activation 0.42 ± 0.04 z-ΔF/F, *p* = 0.0005 for sync-subset. -0.05 ± 0.04 z-ΔF/F, *p* = 0.30 async - subset). These effects were observed regardless of the relative timing of the subset ensemble (Extended Data Fig. 5). This effect cannot be explained by stimulation induced artifacts, as we restrict holographic stimulation to periods around the scan mirror flyback period, where they do not contaminate fluorescent measurements^67,68^. Together, these findings indicate that while all evoked activity recruits recurrent inhibition, synchronous activation can also drive coordinated recurrent excitation. This implies that recurrent excitation and recurrent inhibition integrate differently, with recurrent inhibition integrating over a longer duration than excitation.

## Synchronous neural ensemble activation suppresses neural circuit activity

Finally, we hypothesized that synchronous stimulation may selectively drive activation of certain functionally defined neurons, which in turn drives the behavior. We examined the response in non-targeted cells during sync and async stimulation. To avoid possible contamination by off-target light, we excluded cells nearby any targeted cell^63,67,68^, and to remove the direct effects of evoked movements we excluded all trials that evoked movement. Consistent with previous results^68^, we found that most non-targeted cells reduced their activity in response to activation of targeted neurons (Fig. 5a-c). We observed no difference in the cells’ suppressed responses between sync and async stimulation at the population level (Sync -0.049±0.006 vs Async -0.052±0.006 z-ΔF/F, p=0.86 Fig. 5d).

**Fig. 5.**
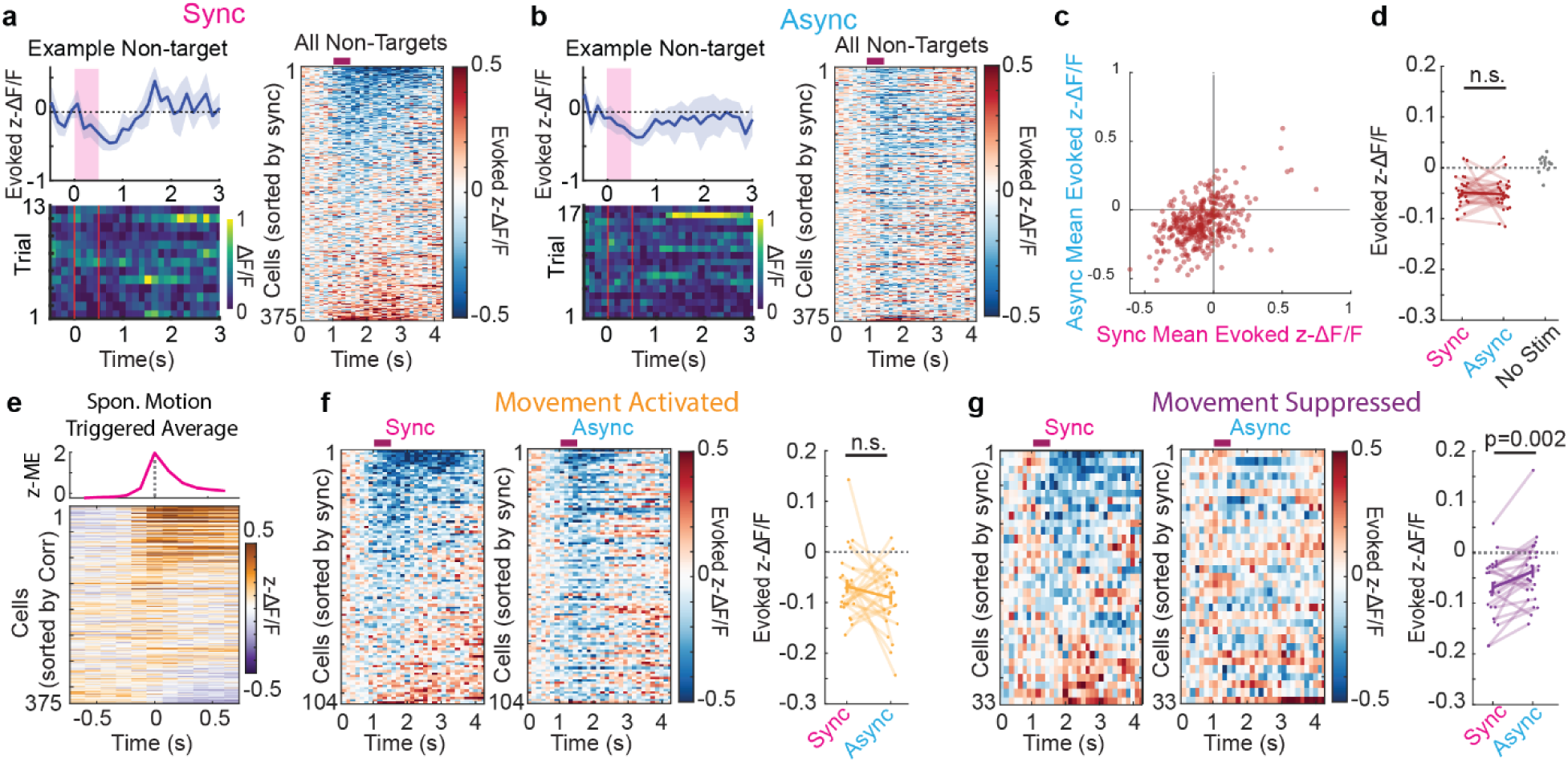
Synchronous and asynchronous activation recruit suppression. a-b. Evoked neural responses of a single example (Left) and all non-targeted neurons (Right) to sync (**a**) and async (b) activation from one example mouse. **c.** Mean of evoked neural response of non-targeted neurons in sync and async conditions. **d.** Mean of evoked neural responses of non-targeted neurons in each ensemble in sync, async, and no-stimulus conditions (Sync -0.049±0.006 vs Async -0.052±0.006 z-ΔF/F, *p*=0.86, signed-rank test). **e.** Example movement (z-ME, top) and neural responses (bottom) in non-stimulation trials aligned to movement onset from one example mouse. **f.** Example evoked neural responses of movement-activated neurons in sync and async conditions from one example mouse. Right: Evoked response of non-targeted neurons in each ensemble in sync and async conditions (Sync -0.07±0.01 vs Async -0.09±0.01 z-ΔF/F, *p*= 0.43, signed-rank test). **g.** Example evoked neural responses of movement-activated neurons in sync and async conditions from one example mouse. Right: Evoked response of non-targeted neurons in each ensemble in sync and async conditions (Sync -0.07±0.01 vs Async -0.04±0.01 z-ΔF/F, p=0.002, signed-rank test).

Next, we categorized non-targeted neurons into movement-activated and movement-suppressed neurons based on their activities during spontaneous body movements (Fig. 1h, 5e). While there was no difference in suppression in movement-activated non-targeted neurons (Sync -0.068±0.01 vs Async -0.09±0.01 z-ΔF/F, p= 0.43. Fig. 5f), we observed significantly more suppression of movement-suppressed non-target neurons in the sync condition (Sync -0.07±0.01 vs Async -0.04±0.01 z-ΔF/F, p=0.002, Fig. 5g). These results indicate that while both synchronous and asynchronous activation recruit local circuit inhibition, they do so in distinct ways targeting different functionally defined cells.

## Discussion

The motor cortex is a critical brain region that drives action planning, refinement and movement generation. However, while much is known about how this region responds during behaviors, limitations in causal testing have obscured how neuronal firing directly evokes movements. We find that the precise activation of small ensembles of excitatory neurons (50-75 neurons) in L2/3 of the naïve mouse motor cortex can directly evoke paw movements (Fig. 1). The efficacy of a given stimulation to evoke a movement is a function of the size of the stimulated ensemble, the ongoing brain state, and, critically, the between-neuron temporal structure (i.e. synchrony).

Several groups have used multiphoton optogenetic approaches to answer variants of the question ‘how many neurons are needed for a behavior?’ in various regions, including visual cortex^53,79^, somatosensory cortex^47^, hippocampus^80^, and, recently, motor cortex^81^. Each of these studies relied on first training their animals to perform a task. While these studies add critical insight, there remains the possibility that optogenetic stimulation triggers a learned cortical activity sequence that the animal is poised to make to drive a trained behavior, possibly through pattern completion^53,82^. In contrast, we focus on naive mice and their spontaneous movements to ensure that the activity we analyze is the product of innate connectivity and not a result of training. We conclude that 50-75 neurons are required to evoke a movement. This is higher than the several dozen neurons reported for behaviors in other studies, presumably as those evoked preplanned behaviors. Simultaneous multiphoton activation of 75 cells is an uncommonly large number of simultaneously activated cells, only being achieved in several, mostly proof of principle, papers^62,83–85^. We show that both the number of neurons driven and the degree of activation of that group affects the efficacy of evoking a movement.

We find that the way a neuron fires during a spontaneous movement is a poor predictor of whether that neuron will drive an *evoked* movement (Fig. 1h-i, 2h). In other words, we find that ‘spontaneous movement’ cells in the motor cortex do not necessarily drive movements. This finding is reminiscent of recent results with visual cortex stimulation^79^. While this observation is counter-intuitive, it reinforces the model that the primary role of superficial motor cortex is not strictly movement generation but rather the processing of movement information^86^. In contrast, the way a targeted neuron fires with respect to the population activity, was a better predictor of that neuron’s effect on movement (Fig. 1j). These more highly coupled cells may represent a subset of cells that are more densely connected to their neighbors^48,77^.

Neural population activity in awake animals can take on many different population activity states regulated by attention and neuromodulation^87^. We developed a computational model that identified a ‘permissive state’ that motor cortex neurons must inhabit before stimulation in order to evoke a movement (Fig. 2). This state is distinct from the premotor state, and most highly weights ongoing activity of non-targeted cells. We find the amount of time a mouse spends in this permissive state is a better predictor of whether that ensemble will evoke a movement than the choice of which neurons are targeted (R^2^ = 0.88 permissive state vs R^2^ = 0.37 population coupling). Together this highlights the importance of population activity in the regulation of motor behavior.

A major emphasis of this study was to understand the relative contributions of spike number and spike synchrony, i.e. are motor cortex evoked behaviors controlled by ‘rate codes’ or ‘timing codes’^14^. While there is evidence both for^35,36^ and against^37,38^ timing codes, the most common multidimensional analyses of neuronal activity in motor cortex tend to use an underlying rate-code assumption, with neural activity binned in tens-of-milliseconds bins (e.g., 10-50ms)^88–94^. Our data shows that temporal offset of 5ms can have a profound effect on the effectiveness of stimulation, with asynchronous activation abolishing our ability to drive movements (Fig. 3). While this finding is agnostic to the contribution of ‘rate codes’ in the motor cortex, it argues strongly that ‘timing codes’ must be potent.

Consistent with previous results^68^ we find that activating more cells drives increasing amounts of network suppression, and this suppression was nearly equivalent between the synchronous and asynchronous stimuli (Fig. 4). However, surprisingly the synchronous stimulation also caused an apparent increase in recurrent excitation, as seen by the faster rise time in the directly stimulated cells. Our controls indicate that this cannot be explained by a difference in cell illumination timing or strength (Extended data Fig. 4 and 5). This implies that the mechanism of action of cortically evoked movement is via recurrent excitation of synchronous activation driving faster responses, with recurrent excitation being more sensitive to timing than inhibition.

Finally, while non-targeted cells are mostly suppressed by both synchronous and asynchronous stimulation, certain classes of neurons appear especially sensitive to synchronous stimulation (Fig. 5). Cells activated by spontaneous movements show a heterogeneous response, whereas cells suppressed by spontaneous movements are less affected by asynchronous stimulation even in trials that did not evoke movements. We conclude that these classes of cells are selectively sensitive to the timing rules of cortical activation.

Taken together we show that direct stimulation of Layer 2/3 motor cortex can evoke movements. The chosen cells, state of ongoing activity and, critically, temporal pattern of activation all play a role in gating whether a stimulus will evoke a movement. We provide direct evidence for the importance of timing codes in motor cortex behavior, demonstrating that small timing differences play a critical role in the effect of neural activity.

## Methods

### Animals

All experiments were performed with approval of the Institutional Animal Care and Use Committee (IACUC) of Rutgers, The State University of New Jersey (Protocol number; #PROTO202200005). Male and female EMX1-Cre mice (Jax stock #005628) were used for all experiments except measuring physiological point spread function (PPSF) (Fig. 1b). To measure PPSF, tetO-GCaMP6s (Jax stock #024742) mice crossed with Camk2a-tTA mice (Jax stock #003010) were used. Animals between 6 weeks, at age of surgery, to 6 months were used. Animals were group-housed in cohorts of five or fewer on a reversed 12-hour dark / light cycle at an ambient temperature and humidity and had free access to food and water. Experiments were performed during the dark phase.

### Surgical procedures

Mice were anesthetized by isoflurane (5% induction, then 1.5-2.0 % maintenance) through oxygen-enriched air. Mice were secured in a stereotaxic frame (Model 1920, Kopf) and kept warm with a heating pad (Rodent Warmer X1, Stoelting). Their eyes were covered by a protective eye lubricant during the surgery. Extended-Release Buprenorphine (Ethiqa XR, 3.25 mg/kg) was subcutaneously injected for analgesia, and bupivacaine (3 mg/kg) was injected under the skin over the skull for local anesthesia. Dexamethasone (2 mg/kg) was subcutaneously injected to reduce cerebral edema during craniotomy. Anesthesia level was monitored via breathing rates and foot and tail reflexes before and during surgery.

A small incision along the midline was made on the scalp, then the scalp over the skull was carefully cut and removed. The periosteum was removed using sterilized cotton swabs, and the skull was lightly etched with a sterilized needle. A custom-designed titanium headplate was implanted on the skull. The headplate was initially secured with Vetbond (3M), then permanently secured with C&B Metabond (Parkell). A craniotomy was made over a left caudal forelimb area (CFA^42,76^, 1.5 mm lateral, 0.5 mm anterior from bregma) for motor cortex stimulation or over a left primary visual cortex (V1, 3.0 mm lateral, 1.0 mm anterior from lambda) using a hand drill (503599, World Precision Instrument) with a burr drill bit (200 - 400 µm diameter, Stoelting). To prevent the brain from heating by drilling, sterilized cooled phosphate buffer saline (PBS) was occasionally applied on the skull. During and after craniotomy, bleeding was controlled by sterilized PBS and SurgiFoam (Ethicon).

For the injection of adeno-associated virus (AAV), a pulled and beveled glass pipette (tip diameter, approximately 25 um) (2000-001, Drummond Scientific) was first back-filled with mineral oil then front-filled with AAV vector. The pipette was slowly lowered into the brain with the help of a motorized manipulator (MP-225A, Sutter Instrument). For EMX1-Cre mice, AAV-PHP.eb-CAG-DIO-stChroME2s-mRuby3 (Addgene, 170163, AAV preparation done by Virovek, ca 50-250 nl, diluted by sterile PBS, 2.5 - 5.0 E12 vg/ml) and AAV2/9-syn-FLEX-jGCaMP8m^95^ (Addgene, 162378-AAV9, ca 50-250 nl, diluted by sterile PBS, 1.5 - 6.0 E12 vg/ml) were mixed and injected into the CFA or V1 approximately 350 um below the dura using a micro injection syringe pump (UMP3 with MEH3SF10, World Precision Instrument). For tetO-GCaMP6s mice crossed Camk2a-tTA mice, AAV-PHP.eb-TRE-stChroME2s-mRuby3 (Addgene, 170175, AAV preparation done by Virovek, ca 100-300 nl, diluted by sterile PBS, 1.3 - 2.5 E12 vg/ml) was injected into the CFA. The glass pipette was kept in place 3-5 minutes after the AAV injection to ensure adequate spread and prevent backflow of AAV during retraction of the pipette. A cranial window was constructed by adhering two 2.0 mm diameter circular coverslips (Thickness No. 1, Matsunami US) or one 2.0 mm diameter circular coverslip (Thickness No. 3, Matsunami US) to the bottom of a 3.0 mm diameter coverslip (Thickness No. 1, Matsunami US or CG10E1, Thorlab), using optical adhesive (NOA81, Thorlab). The coverslip was placed on the craniotomy and secured in place with C&B Metabond (Parkell). Following the surgery, mice were placed into a heated recovery chamber until they became awake and started to walk.

### Two-photon holographic microscope system

*In vivo* two-photon calcium imaging and two-photon holographic optogenetic stimulation was performed using a movable-objective microscope system (MOM, Sutter Instrument) combined with a 3D-SHOT system^65,67,68^. The microscope was equipped with a resonant-galvo scanning system (Sutter Instrument) and a pulsed Ti:sapphire laser (λ = 920 nm, Discovery NX, Coherent). A motorized three-axis system (MP-285, Sutter Instrument) connected with a microscope head enabled locating an objective lens above targeted brain area. An electrical tunable lens (EL-10-30-TC-NIR-12D with ICC-4C-500, Optotune) was placed immediately before the scanning system in the imaging pathway for fast multiplane volumetric imaging. A polarizing beam-splitter (PBS, PBS255, Thorlab) merges the imaging laser and holographic stimulation laser paths. The microscope system was controlled by ScanImage software (MBF Bioscience). Green and red fluorescent photons were collected with an objective lens (20x XLUMPLFLN Objective, 1.0 NA, Olympus). Photons were separated by a dichroic mirror (T565lpxr, Chroma) and barrier filters (green: ET525/70m-2p, red: ET605/70m-2p, Chroma), and measured by PMTs (H10770PA-40-04, Hamamatsu photonics). The imaging frame was 512 x 512 pixels, and the frame rate of multiplane imaging was approximately 7.5 Hz per plane for three-plane imaging and approximately 5.6 Hz for four plane imaging. Field of view (FOV) of two-photon images were approximately 800 µm x 800 µm. To minimize the excitation of opsin by scanning of the imaging laser (i.e., cross-talk) ^62^, we restricted imaging laser power less than 70 mW^67^.

Two-photon optogenetic stimulation was performed using a custom-designed 3D-SHOT system. The details of the 3D-SHOT system have been described previously^63,65,67,68^. Briefly, stimulation used a 1030nm pulsed laser (λ = 1030 nm, 50W, aeroPULSE FS50, NKT Photonics) at low repetition rate (approximately 1.25 MHz) and a fast spatial light modulator (HSP1k SLM or HSP1920HB SLM, Meadowlark Optics) to direct light to different targets. Phase masks are generated using custom-written MATLAB code using the Gerchberg-Saxton algorithm^96^. The stimulation laser was gated^67,68^ such that the stimulation artifact only occurred on the edges of the imaging field of view to prevent stimulation artifacts from contaminated neurophysiological signals.

### Setting a custom power per cell

To compensate for the variable intrinsic excitability and the level of opsin expression across neurons, we determined an individual power to drive each targeted neurons^63,67,68^. Briefly, in each experiment, we stimulated groups of neurons with a train of five 5ms pulses at various intensities. The calcium responses were analyzed online using either custom-written MATLAB codes or an online interpretation of the CaImAn onACID algorithm^97,98^ adapted for MATLAB^68^. The target power used (28 ± 12 mW (mean±SD) for each cell during the rest of the experiment was set as the minimal power necessary to drive that cell to the saturating level. Neurons that were not reliably activated during this test were not used as ensemble members. As the optimal power is unable to be determined for opsin-free control experiments, stimulation powers comparable to other experiments were manually assigned to selected neural ensembles.

### Recording of physiological point spread functions

To measure the resolution of our holographic targeting, we measured the physiological point spread functions (PPSFs)^63,68^ for a total of 40 cells (N = 40 PPSFs, 4 FOVs, 2 mice). Holograms were offset radially in 3 µm steps (range: -3 to 30 µm) and axially in 6 µm steps (range: -6 to 60 µm) from the target neuron. The resulting calcium activity (ΔF/F) is fit with a Gaussian, and the full-width half max (FWHM) was calculated.

### Neural ensemble stimulation

Photoactivatable neurons were randomly assigned into ensembles and driven while observing for evoked body movements. Groups of 25, 50 and 75 photoactivatable neurons were stimulated at 30 Hz with 1, 5 or 15 pulses (Motor Cortex stimulation varying number of cells, N = 71 ensembles, 29 FOVs, 10 mice; varying number of pulses, N = 46 ensembles, 19 FOVs, 9 mice. Visual Cortex control experiments, N = 15 ensembles, 6 FOVs, 3 mice. Motor Cortex without opsin control, N = 24 ensembles, 8 FOVs, 3 mice). Each instance of laser stimulation consisted of a 5 ms pulse. The order of stimulation sequence was pseudo-randomized.

### Synchronous and asynchronous neural ensemble stimulation

Ensembles of 50, 70 or 75 photoactivatable neurons were selected, and stimulated in two different patterns (N = 25 ensembles, 11 FOVs, 5 mice for each matched pattern). For synchronous stimulation, all members of an ensembles were simultaneously stimulated with 15 pulses at 30 Hz (5 ms pulse width). For asynchronous stimulation, the same ensembles were split into three roughly equal size sub-ensembles (25, 25, 25 targets; 16, 17, 17 targets, or 23, 23, 24 targets) and stimulated in an interleaved manner with a 5 ms gap between the offset of the previous stimulation pulse and the onset of the next stimulation pulse. 22 ensembles used the HSP1k SLM and 3 used the HSP1920 SLM (both Meadowlark), both SLMs can fully change targets by 5ms. Thus, for both synchronous and asynchronous conditions, the selected neurons were stimulated with the same parameters including frequency, and the number of pulses, but not intercell timing.

### Subset stimulation experiments

In these experiments, targeted ensembles of either 50 or 75 total cells were stimulated in three different patterns: synchronous, asynchronous or subset stimulation. Synchronous and asynchronous stimulations were performed as described above. In the subset stimulation groups, only one sub-ensemble of the three parts of the asynchronous condition were driven. The timing of each subset stimulation was matched to where that sub-ensemble would have been driven during the asynchronous trial type with respect to trial onset. In all conditions, the targeted neurons were stimulated at a frequency of 30 Hz with 15 pulses (3 ms pulse width). N = 4 starting ensembles, split into 12 total single-subset partitions, 4 FOVs, 2 mice. During subset stimulation, although only one sub-ensemble was stimulated, the SLM continued to display multiple holograms as in the async condition.

### Two-photon Image Processing

Two-photon images were processed using Suite2P^99^. The images were motion-corrected, and regions of interest (ROIs) were automatically generated. Next, experimenters curated ROIs and sorted them as neurons or not. A neuropil signal was calculated for each ROI via Suite2p, and each ROI signal was subtracted from the neuropil signal multiplied by a correction coefficient (0.7). The resulting signals represent the raw fluorescent calcium trace for individual neurons. Raw fluorescence was converted to ΔF/F, using the 10^th^ percentile of a moving baseline to determine F_0_. The resulting ΔF/F vector was converted to z- ΔF/F via z-scoring of the unrolled fluorescent trace.

### Exclusion Criteria

We excluded the data similarly as described previously^68^. Trials were excluded in case 1) half or more than half of the targeted neurons failed to respond to the holographic stimulation (beyond 0.25 zScore) except experiments of a small number of pulse and no-opsin stimulations; 2) the brain was found to have moved more than 4.8 um (3 pixels) during stimulation; 3) the mouse body movement during approximately 250 ms before holographic stimulation was beyond 95 % confidence interval calculated from all trials within a session (see also Mouse body movement analysis, and Extended Data Fig. 1e); 4) Imaging laser unexpectedly cut off during experiment (total six non stimulation trials). Neurons were excluded in case 1) they were found in an “off-target” region, 15 µm from targeted neurons radially and 30 µm radially from the targeted neurons from one imaging plane away (Figure 5 only); 2) non-targeted neurons were located within a stimulation laser artifact area (All Figures). Neural ensembles were excluded if more than 50% of the targeted neurons were not detected via suite2p process except no-opsin stimulation, or if more than 50% of trials were excluded from that condition.

### Stroboscopic Imaging for SLM High Speed Performance Testing

To measure the rate the SLM can transition phase mask, we performed stroboscopic imaging as previously described^67^. In each round, three single-target holograms are sequentially displayed on a thin fluorescent slide while imaging with a substage camera (CS2100M-USB, ThorLabs), transitioning from Hologram A to B to C and back to A at a frequency of 100Hz using either SLM (HSP1K or HSP1920HB, Meadowlark). In any given trial, a single 0.5 ms ‘stroboscopic’ laser pulse illuminates the slide at any time within the trial duration. The entire hologram sequence is illuminated after 100 or 150 trials, for trial durations of 50 and 75 ms, respectively. Each round of a distinct hologram sequence (N=5 rounds, 50 ms, HSP1k SLM; N=3 rounds, 75ms, HSP1920HB SLM) consists of 10 replicates (1000 or 1500 trials).

Holograms are randomly chosen within the field of view and matched in power efficiency. After measuring the intensity of the holograms, we average the 10 replicates and normalize. To ensure holograms are transitioning at the required experimental stimulation rate, we calculate the average time it takes for a hologram to reach 90% if its illumination peak from the end of the SLM trigger signal by fitting a Hill function. To ensure each experimental ensembles receives equivalent amounts of light, we calculate the area under the curve for 5 ms at timepoints 5, 10, 15, and 30 ms after the SLM trigger signal.

### Fluorescence intensity as a function of hologram size

To measure the variability in delivered power across varying sizes of ensembles^52^, we used the substage camera to measure the intensity of 8 individual ‘test’ targets on a thin fluorescent slide, either alone in a single-target hologram or within a multi-target hologram comprising of 5, 10, 25, 50, or 75 randomly chosen targets spread across three planes. For every test target and hologram size, 5 replicates were performed. Intensity data is then normalized to a single target.

### Mouse body movement analysis

Mouse behavior was monitored during experiments with a CMOS camera (acA800-510um-Monochrome, Basler) equipped with a fixed focal length lens (33-303, Edmund Optics) and a short-pass filter (86106, Edmund Optics) in a filter mount (65800, Edmund Optics). The camera was located on the right and front side of the mouse with an infrared LED (810 nm, Thorlab). Each frame was acquired at 200 Hz. During offline analysis, amplitude of the body movements were quantified using FaceMap^100^. Motion energy calculated from the area covering mouse paws and used as a measure of body movements. The motion energy data was lowpass-filtered and z-scored in each session.

Pre-stimulation movement trials (see exclusions above) were defined as: trials where the average movement in a ‘pre-stimulus’ window, 250ms before stimulus onset, exceeds the 95% confidence interval of all pre-stimulus windows. Exclusions were performed identically in stimulation and non-stimulation trials. After removing the pre-stimulation movement trials, the body movement data was baselined by the average movement in the pre-stimulus window across all trials in each session. Then, mean movement responses in the movement response window, defined as the 500ms after stimulation onset, were calculated for all trials. If the movement responses during the response window was beyond baselined z score = 0.65, this trial was defined as “movement trial”. To test whether holographic stimulation evoked body movement, fisher exact test was performed between stimulation and non-stimulation trials.

### Hand trajectory analysis

During offline video analysis, the paw center was detected and tracked by using markerless pose estimation algorithm, DeepLabCut (DLC)^101^. The mouse’s right paw center was manually labeled for all session data to train the model. Then, video data from every recorded session was analyzed using DLC to detect the paw center in each video frame. Two sessions were excluded from this trajectory analysis, as the paw was not detected in more than 10% of the total frames in those sessions. In all other sessions, the paw was detected in 98 ± 0.02% of frames (mean ± STD) before data exclusions described above. To quantify the similarity of paw movements, we computed the cosine similarity of the tracked paw coordinates. Cosine similarities were computed across three distinct trial conditions within each session: (1) between stimulation trials within the same hologram, (2) between no-stimulation trials (as spontaneous movement similarity), and (3) between stimulation and no-stimulation trials. In stimulation trials, we searched for the paw movement within the time window during the holographic stimulation. To avoid the detection of ongoing movements before stimulation, in addition to pre-stimulus trial exclusion, the first 100 ms after the stimulation onset was not used. In no-stimulation trials, we searched for the paw movement as those spontaneous events. Pre-stimulus movement trials were also used to analyze hand trajectories in no-stimulation trials. Similarly in stimulation trials, the first 100 ms after the trial onset was not used to avoid the detection of ongoing movement. Multiple paw movements in a single trial were analyzed in no-stimulation trials. In each session, if the number of detected paw movements were less than two events, those data were not used for further analysis. The angle of the vector of a movement was calculated relative to the movement’s origin and displayed in polar coordinates.

### Movement cell classification

Cells were classified as movement-active, movement-suppressed or non-movement related cells using neural activity and motion energy data in no-stimulation trials. First, motion energy data was down sampled, followed by a cross-correlation calculation between individual cell activity and motion energy. Cells were classified based on their cross-correlation values and the timing of their peak response. A cell was classified as movement-active if its peak cross-correlation exceeded 0.1, or movement-suppressed if it fell below -0.1, provided the maximum correlation occurred between -1 and 5 frames lags (∼130 ms before to ∼670 ms after movement). Cells with cross-correlation values between -0.1 and 0.1 were classified as non-movement correlated. A motion triggered average was created detecting the onset of bouts of movement activity.

### Calculation of target cell distances

The Euclidian distance was calculated between targeted cells within each ensemble. After calculating the distance of all pairs, the distances were averaged in each ensemble. This mean of the targeted cell distance was used to analyze the relationships between the targeted cell distance and the mean of the stimulation-evoked targeted cell responses (population response) and the movement probability.

### Active cell classification

Similarly to Movement cell activity classification, the concatenated cell activity data and down sampled motion energy data in no-stimulation trials were used for this analysis. Using regression analysis, the movement-explained component was subtracted from the cell activity (residual activity). To determine the event rate, we calculated the fraction of frames in the residual activity that exceeded the activity threshold (*z* = 2.5). Within a session, cells above 75 and below 25 percentiles were classified “Active cells” and “less-Active cells”, respectively. The proportion of Active and less-Active cells within each ensemble were calculated to analyze the relationship between the nature of cell activity and movement probability.

### Calculation of population coupling

We calculated the ‘population coupling’, i.e. the correlation of neural activity in no-stimulation trials between each neuron and all other non-stimulated neurons (population) within a session.

### Non-targeted cell analysis

We analyzed neural responses of the non-targeted cells to synchronous and asynchronous stimulation. As described in exclusion criteria, non-targeted cells located within stimulation laser artifact and off-target region were excluded from analysis. In addition, to test the circuit neural activity without movement effects, trials with evoked movements were excluded. The movement trials were classified as described above (Mouse body movement analysis). Cells were sorted into movement-active and movement-suppressed cells, as described above (Movement cell classification).

### PCA-based permissive state analysis

Motion energy per trial and the classification of trials as high or low movement was determined as described above. For each stimulated ensemble a model was trained to predict the trial-wise movement energy from the neural activity in 5 frames prior to photostimulation. The model chosen for this analysis was a ridge (L2-regularized) regression (alpha = 1), implemented using Python’s scikit-learn package. To avoid biasing models, any experiments in which there were less than 2 evoked movements during stimulation trials were excluded from further analysis and model fitting. The first model used compared trial-wise activity of all neurons, restricted to the first 6 frames of calcium imaging data, to the evoked motion energy. A weight was fit to each neuron individually. To evaluate the model for overfitting, trial-wise motion energy values were shuffled a thousand times, such that within each iteration a model was trained on the shuffled motion energy values and used to predict the shuffled motion energy values. The second ensemble-based permissive state analysis followed the same method as the first but first reduced the dimensionality of neural activity using PCA. Principal components were found using singular-value decomposition of a flattened matrix consisting of all neurons’ response across all non-stimulation trials. Subsequently, for each trial, the neural response from both stimulated and non-stimulated trials was centered and projected onto the first four components of this principal component space. Model fitting and evaluation followed the same procedure as above. Using the weights and intercept from the fit model, neural activity of non-stimulated trials was converted into predicted motion energy.

### Permissive state analysis based on functionally defined neuronal groups

For every neuron, evoked movement correlation was determined by comparing the mean neuronal activity in the 5 imaging frames immediately following stimulus onset to the post-stimulus movement energy across all trials. The top 5% most correlated neurons were categorized as ‘positively correlated,’ the bottom 5% most negatively correlated neurons were categorized as ‘negatively correlated,’ and the remaining neurons were categorized as ‘not correlated’ or ‘null.’ This procedure was applied separately to targeted and non-targeted neurons. For the shuffled control shown in Fig. 2c, 200 linear regression models were trained to predict different permutations of every session’s ME values. Separately, another form of shuffled control was performed in Extended Data Fig. 3g,k where ME values per trial were shuffled within each session before categorizing neurons based on their movement correlation. This permutation of ME values (and neuronal groups) was repeated 200 times, and the mean explained variance across shuffles is shown for each ensemble. Statistical comparison of each model to its shuffled control for each session use paired t-tests, and a Bonferroni correction was applied as necessary for multiple comparisons.

### Linear and logistic regression modeling and cross validation

Ridge (L2-norm, alpha=10) linear regression was used to predict the values of mean motion energy in the post-stimulation window for each trial across all sessions from the cell-averaged neural activity of the above-described six groups of neurons. To test model generalization, either 20% of trials were held out for each session, or 20% of sessions were held out in a random 5-fold cross validation procedure, and then the models were then tested on the held-out data. Explained variance was calculated by from the correlation between predicted and measured ME Statistics were performed on Pearsons R^2^ computed for each session, averaged across the 5 cross-validations. Some sessions were withheld from analyses if 20% of held-out trials amounted to less than two trials. For each session, a logistic regression was then performed to predict whether a stimulated trial was classified as high or low motion energy as described above. In Fig. 2j, the predicted fraction of high ME trials was confidence weighted by the dynamic range (max – min) of motion probabilities predicted by the logistic classifier.

### Brain state metrics

To estimate the time spent in the permissive state during non-stimulated trials, each trial was split into six 5-frame long time windows, and the neural activities for the six neural groups described were averaged within each window. The linear and logistic models described above (trained on pre-stimulus neural features to predict evoked movements on stim trials) were applied to these neural features, resulting in a both a continuous valued prediction of movement probability, and a binary prediction of high or low movement energy for each window. Fig. 2k displays the continuous movement probability versus time for an example trial, and Fig. 2l displays the fraction of non-stimulated time windows classified as permissive versus the measured fraction of stim trials with large movements.

### Parameter importance analysis

To understand which of the six neural features were important for successful motion energy prediction, we trained six additional models were one featured was removed. The change in explained variance (ΔR^2^) for each partial model was computed relative to the full model trained on all six features. One-sample t-tests with Bonferroni correction for multiple comparisons were used for statistical analyses. For examination of model weights, statistics were performed on the values of weights from the five models trained on 80% of trials during cross-validation as described above.

### Spontaneous motion-triggered average neural data analysis

For every non-stimulated trial, a convolutional filter was used to find the largest magnitude spontaneous movement. Neural activity and ME were then aligned to 2 imaging frames prior to motion onset for comparison to stimulation trials were motion typically occurs within 2 frames of stimulation. When spontaneous motion was used to define neuronal groups, movement correlation was determined based on the correlation between the average neural data in the 5 frames before the movement and the motion energy after movement onset.

### Statistical analysis

Statistical analysis was performed using MATLAB (MathWorks). Alpha level was set at 0.05. Fisher exact test, Wilcoxon signed rank test, Wilcoxon-Mann-Whitney test and Kruskal Wallis test were performed. Data are presented as mean ± SEM unless otherwise stated.

## Data availability

Source data to generate figures will be provided upon publication.

## Code availability

### The code used for analysis and figure generation is available at

Github - OldenburgLab- Public

## Acknowledgements

The authors wish to thank Alessandro Galloni for preliminary data analysis, Hailey Thomas for experimental support and James Whitley for his helpful discussions on an earlier version of this manuscript. This work is supported by the Brain & Behavior Research Foundation Young Investigator Grant to M.H.; CSCR26FEL012 from the NJ Commission on Spinal Cord Research to B.G.; funds from Rutgers Biomedical and Health Sciences, and NIH RF1MH135576 to A.D.M; and the Whitehall Foundation, the Searle Scholars Program, NIH R00EY029758, and DP2MH140135 to I.A.O.

## Author Contributions

M.H. and I.A.O designed all experiments. M.H. Performed all the all-optical experiments. B.G. performed the stroboscopic, PPSF, and intensity balancing experiments as well as calibrated the microscope. I.A.O. and B.G. built the holographic optogenetic microscope system. M.S.F. and A.D.M. designed and developed the permissive state analysis based on functionally defined neuronal groups. V.J.A. developed the ensemble-by-ensemble permissive state analysis. All authors wrote the manuscript. I.A.O supervised the work.

---

**Extended Data Fig. 1.**
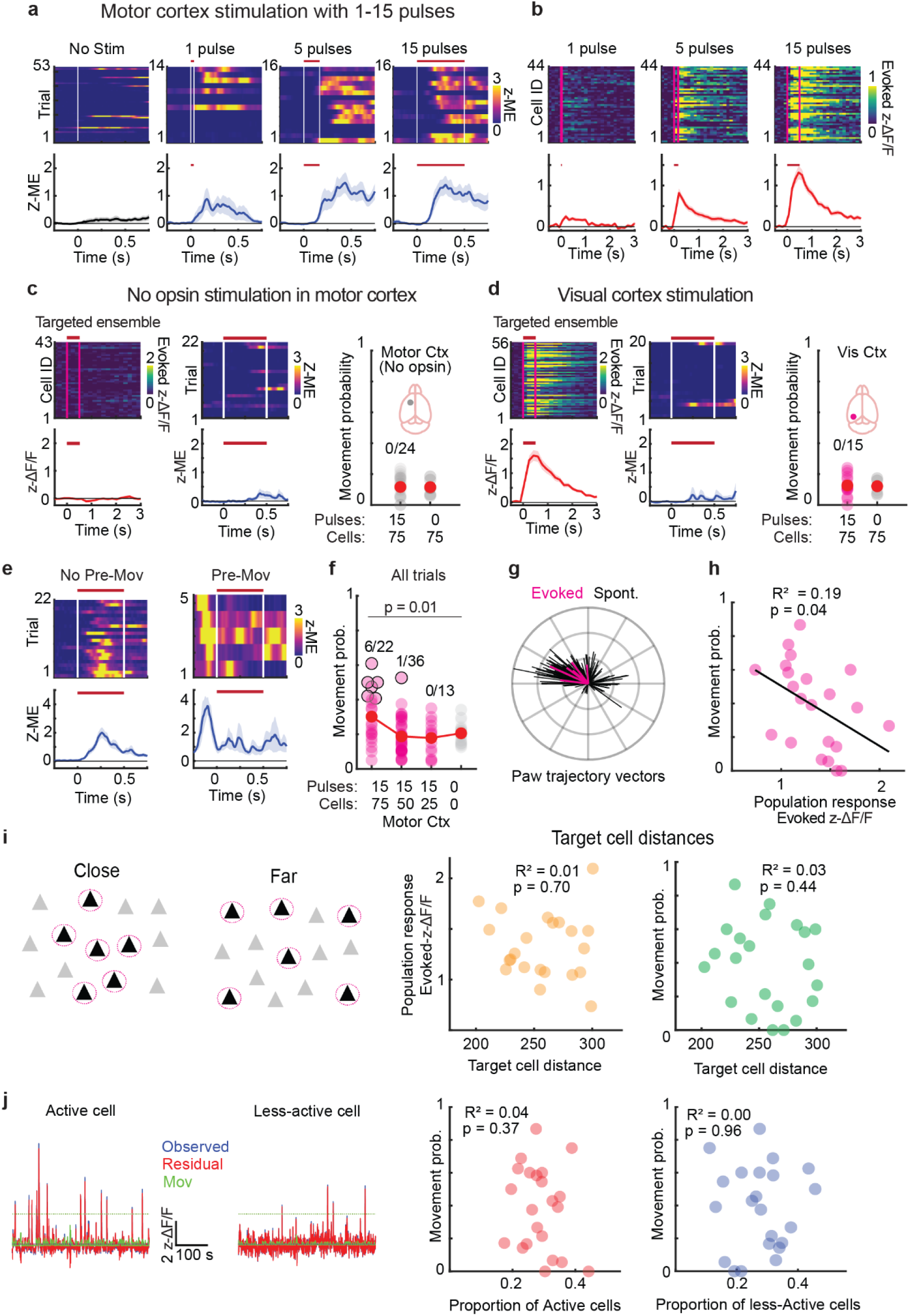
Two-photon holographic stimulation with different conditions. **a**. Evoked body movements with the different number of stimulation pulses (1-15 pulses, 75 cells) from one example mouse. **b**. Evoked neural responses of the targeted neurons with varying numbers of stimulation pulses from the same mouse shown in a. **c**. Evoked neural responses (Left) and body movements (Middle) of the motor cortex stimulation without opsins from one example mouse. Right: Movement probability across all ensembles. Motor cortex stimulation without opsins did not evoke body movements (Fisher exact test, 0/24 movements-evoked ensembles). **d**. Evoked neural responses (Left) and body movements (Middle) of visual cortex stimulation from one example mouse. Right: Movement probability across all ensembles. Visual cortex stimulation did not evoke body movements (Fisher exact test, 0/15 movements-evoked ensembles). **e**. Body movement data with and without pre-stimulus movement trials from one example mouse. For visualization, the z-score of body movements was calculated based on the 30^th^ percentile data point for this panel. **f**. Movement probability calculated from all trials including pre-stimulus movement trials. **g**. Paw trajectory vectors generated from the paw trajectories shown in Fig. 1g. **h**. The movement probability as a function of stimulation-evoked population responses. **i**. Left: Schematics of the targeted cell distances (close and far conditions). Right: Evoked neural response of the targeted ensembles as a function of targeted cell distance, and movement probability as a function of targeted cell distance. **j**. Left: Example activities of active and less-active neurons in no-stimulation trials from one example mouse. All no-stimulation trials are concatenated within a session. The neural activity of each neuron was separated into observed, movement component, and residual component. Right: Movement probability as a function of proportion of the Active Cell and less-Active cells in the targeted ensembles. In figure panels, red or white vertical lines on heatmaps represent stimulation onset and offset. Opaque magenta rectangles and red horizontal lines also indicate the time window of stimulation. Traces shown in a-e are mean ± s.e.m.

**Extended Data Fig. 2.**
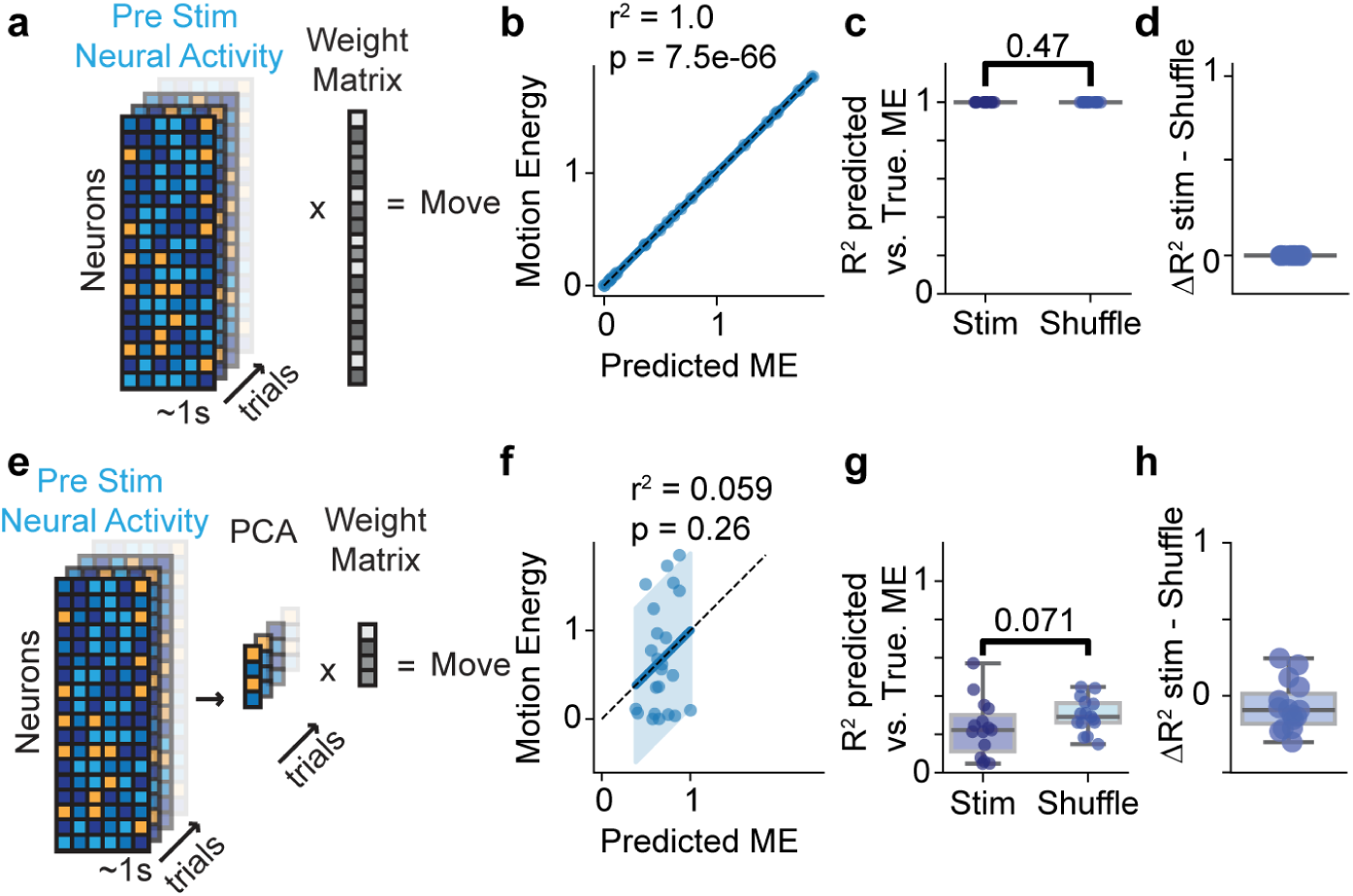
Ensemble by ensemble Permissive state model. **a.** Schematic of ridge regression model using all neurons across stimulation trials. **b.** For an example experiment, correlation between predicted evoked motion energy vs. actual evoked motion energy from stimulation (n = 15 experiments, R^2^=1.0, p=7.5e-66, Pearson). **c.** Difference in correlation coefficients (R^2^) between predicted evoked motion energy vs. actual motion energy for actual stimulation trials vs. the average of a thousand shuffles (R^2^ model: 1.0 ± 0.0, shuffle: 1.0 ± 0.0, p=0.59, rank-sums test) **d.** Difference between R^2^ and average of shuffle accuracies for all-neuron model (ΔR^2^ = 1.7e-6 ± 7.1e-7) **e.** Schematic of ridge regression model using the first four principal components, **f**. as in **b** but for model using first four principal components (R^2^=0.34,p=0.0017, person), **g.** as in **c** but for principal component model (model R^2^: 0.23 ± 0.04, shuffle: 0.30 ±0.02, p=0.0054, rank sum test). **h.** as in **d** but for principal component model (ΔR^2^ = -0.07 ± 0.04)

**Extended Data Fig. 3.**
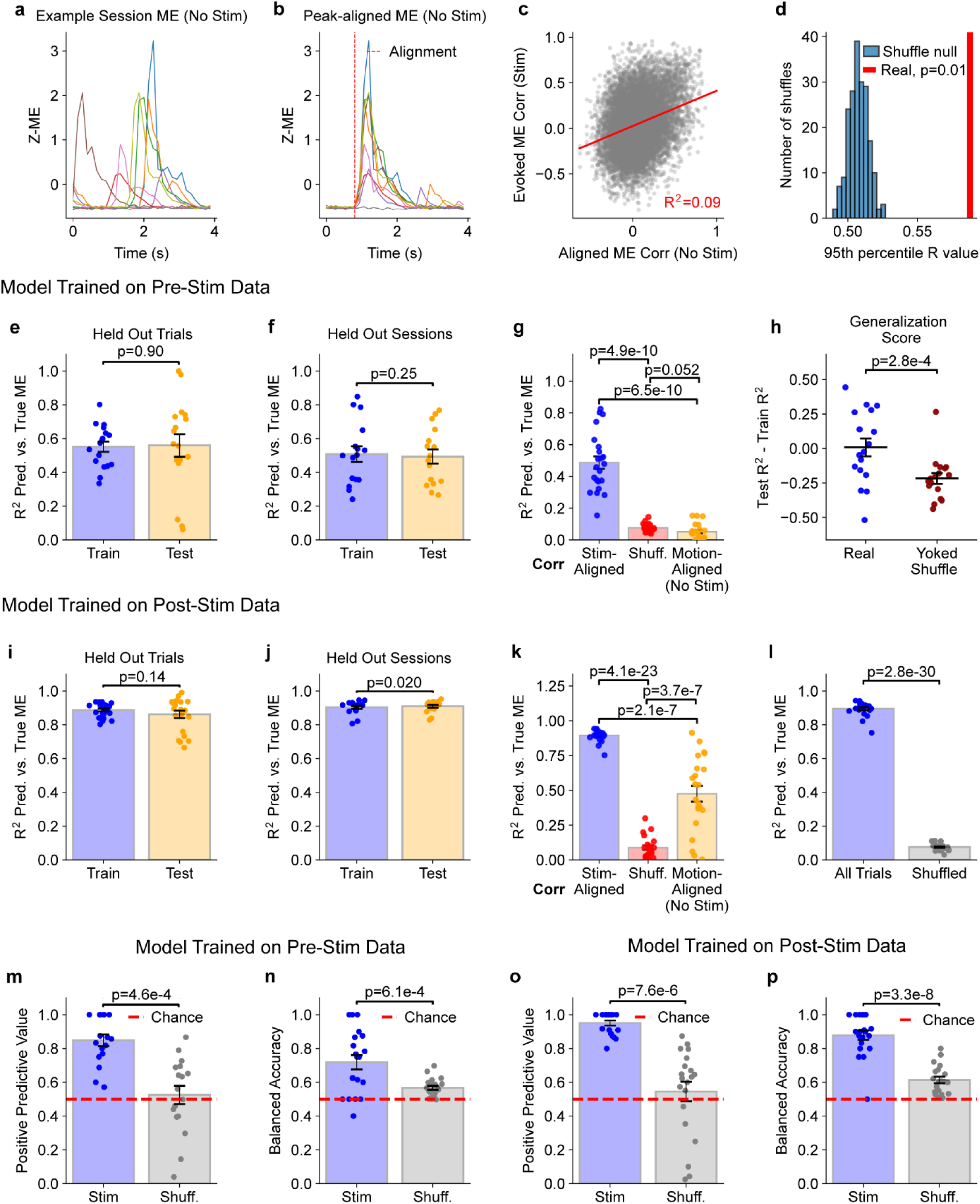
The validation of the permissive state model. **a.** Z-scored motion energy (ME) from non-stimulated trials of an example session. **b**. ME aligned to the largest spontaneous motion using the same trials as **a. c.** After alignment to spontaneous motion onset as in **b**, for each recorded neuron, the average neural activity within a 5-frame window after movement onset was correlated with ME during non-stimulated trials (Pearson, y-axis). Here this is compared to each neuron’s correlation with evoked ME during stimulated trials (Pearson, x-axis). **d**. In Fig. 2, non-targeted neurons were categorized as positively correlated with movement if they were in the top 5% most-correlated neurons in a recording session. Here we compare the 95th percentile correlation (Pearsons) threshold value obtained from actual data to the distribution of threshold values obtained across 200 different shuffles of the ME data across trials within each session. Data shown are averaged across sessions. **e**. Models were trained with 5 different permutations with 20% of trials held out per session, and then tested on the held-out trials. **f**. Models were trained with 5 different permutations with 20% of sessions held out, and then tested on the held-out sessions. In **e** and **f**, data shown are averaged across sessions and permutations, and error represents SEM across the 5 permutations. **g**. Model explained variance is compared to various shuffle controls. In Fig. 2, neuronal feature groups were defined by their correlation with evoked ME calculated from stimulation trials (shown here for reference in blue). A shuffle control first shuffled the ME values across trials within each session before calculating movement correlation and selecting neuronal feature groups (red, average of 200 shuffles). Another alternative model calculated movement correlations from spontaneous movements during non-stimulated trials (orange). **h.** Generalization score is defined as the difference between the explained variance (R^2^) obtained from the train and test phases of 5-fold cross-validation with held-out trials, as in **e**. Here the model in **e** (shown here in blue) is compared to a yoked control model where shuffled ME values were used for all of the following: 1) calculating movement correlations, 2) training, and 3) testing. This yoked model, shown as an average of 200 shuffles, is less able to explain variance of held-out data (lower generalization score). **i-k**. Same as **e-g** for a model trained on neural data from the 5 imaging frames after stimulus onset. **l**. Same as Fig. 2c for the model trained on post-stimulus neural data. **m-n**. Positive predictive value (fraction of positive predictions that are correct) and balanced accuracy (average of true positive rate and true negative rate) for the model in Fig. 2j-l trained on pre-stimulus neural data. **o-p**. Same as **m-n** for the model trained on post-stimulus neural data.

**Extended Data Fig. 4.**
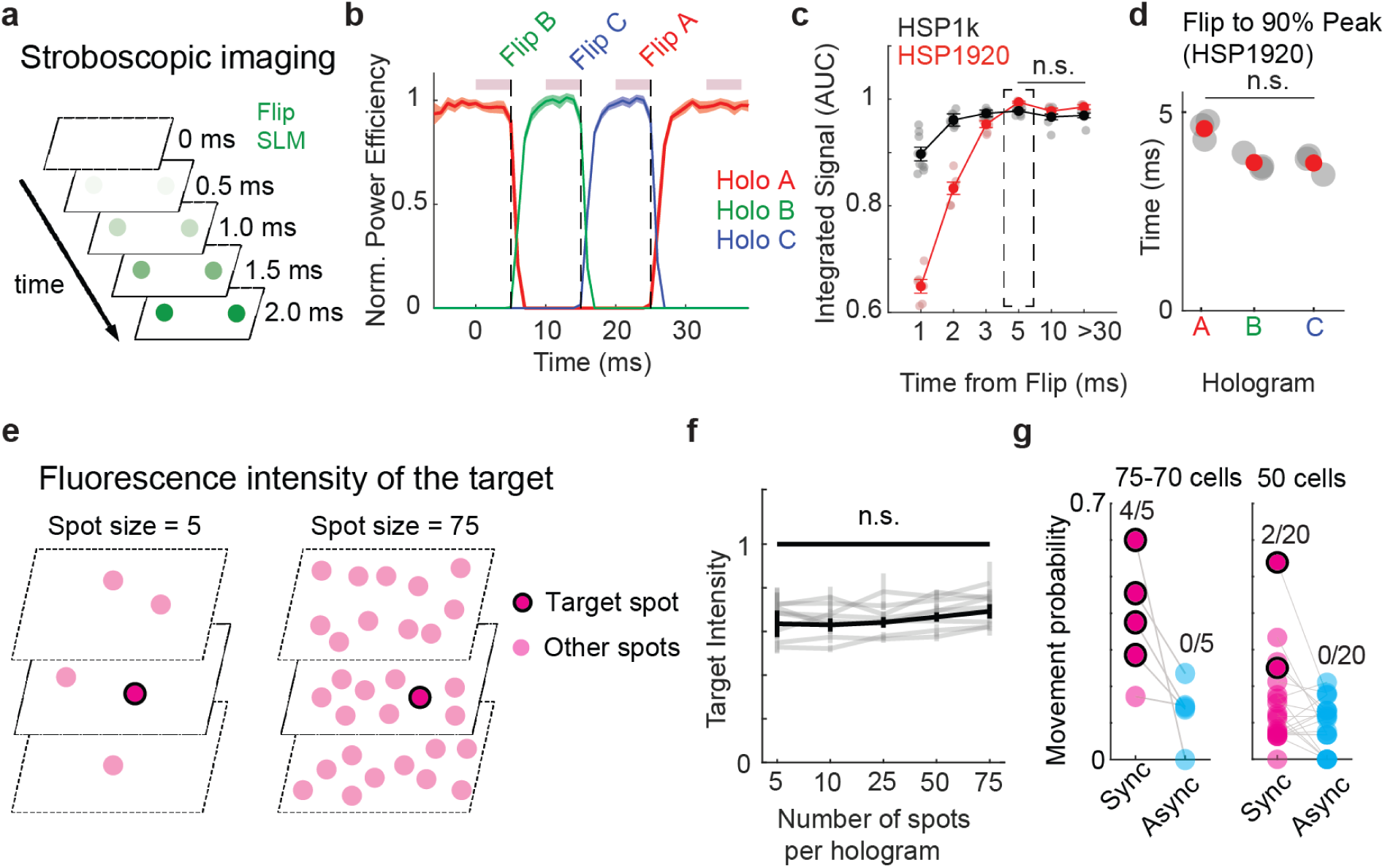
The validation of stimulation efficacy in asynchronous stimulation. a. Schematics of stroboscopic imaging (see methods). b. Power in each of three targets during transition through three holograms as in asynchronous experiment. Each color bar represents the normalized power in a target corresponding to one hologram. Dashed line indicates the flip to the next hologram. Maroon bar indicates when stimulation would occur in an Async experiment. c. Fluorescence intensity of area under the curve. Irrespective of SLM model, AUCs were not different at time points more than 5ms after flipping (p = 0.53, Kruskal-Wallis test, non-significant). The condition used for Async experiments, indicated by dashed box. d. flip time to 90% peak for the HSP 1920 SLM (p = 0.07, Kruskal-Wallis test, non-significant). e. Schematic of power intensity test. Power in a target was compared with a varying number of simultaneously displayed targets. f. Summary of the target intensity across different number of spots. No significant difference between target intensities with various ensemble sizes (p=0.48, Kruskal-Wallis test, non-significant). g. The movement probability of 75 and 50 cells ensemble from the same data used for the Fig. 3d. Traces shown in b and dots and error bars shown in c are mean ± s.e.m.

**Extended Data Fig. 5.**
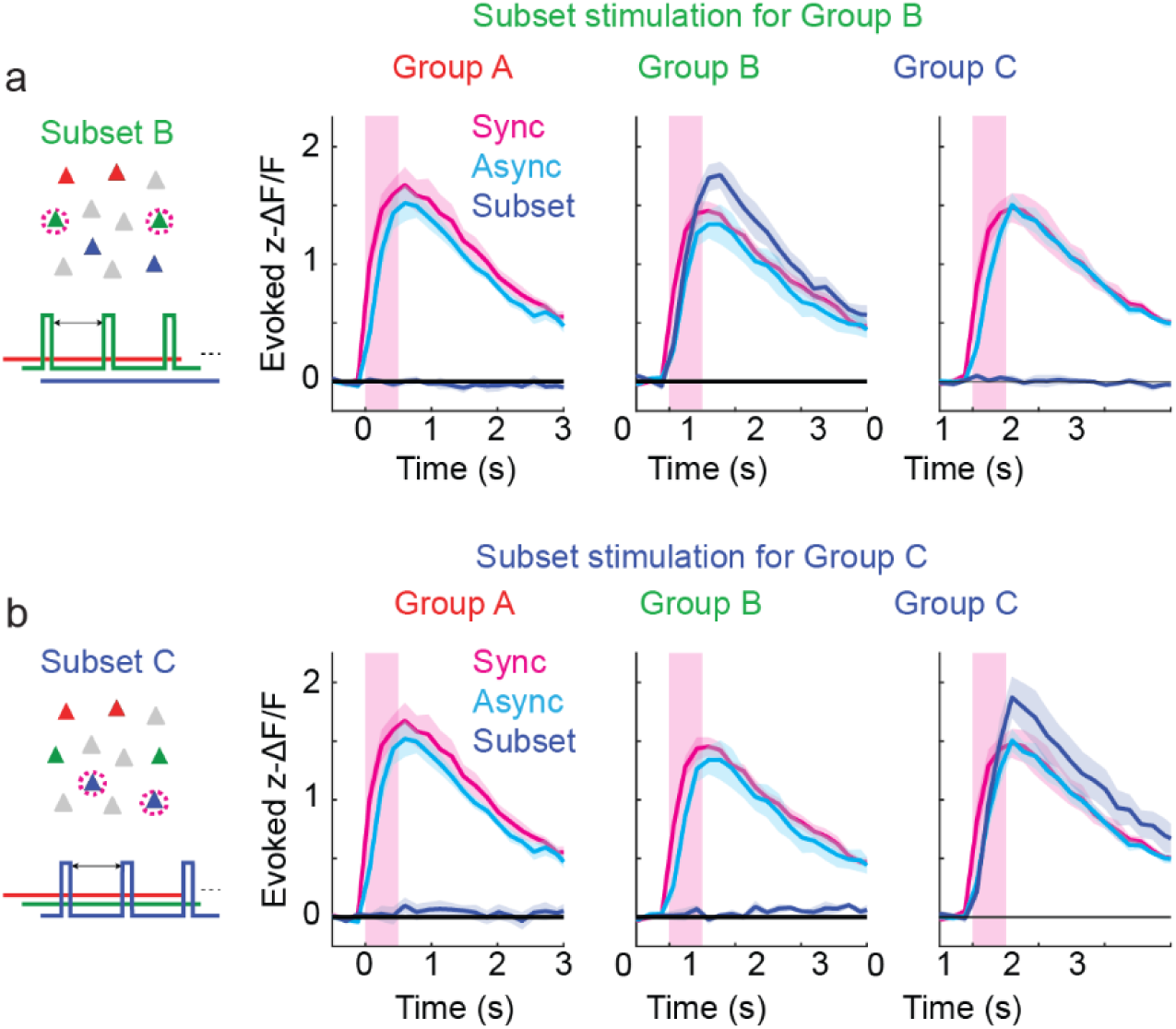
Evoked neural response of single-subset stimulation. **a**. As in Figure 4, Subset stimulation for Group B. Responses of neurons in group A, B, and C during Sync, Async, Individual Subset stimulation of Group B. **b**. As in a but for subset stimulation of group C. Traces are mean ± s.e.m.

